# Poly(ADP-ribose) mediates bioenergetic defects and redox imbalance in neurons following oxygen and glucose deprivation

**DOI:** 10.1101/2023.05.30.542910

**Authors:** M Iqbal Hossain, Jun Hee Lee, Jean-Philippe Gagné, Junaid Khan, Guy G Poirier, Peter H King, Valina L Dawson, Ted M Dawson, Shaida A Andrabi

**Author notes:** Corresponding Author: Shaida A. Andrabi, PhD, Dept. of Pharmacology and Toxicology, 1670 University Blvd., Volker Hall Room 242, University of Alabama at Birmingham, Birmingham, AL 35294, USA. Authors contributed equally (M Iqbal Hossain, Jun Hee Lee).

## Abstract

PARP-1 over-activation results in cell death via excessive PAR generation in different cell types, including neurons following brain ischemia. Glycolysis, mitochondrial function, and redox balance are key cellular processes altered in brain ischemia. Studies show that PAR generated after PARP-1 over-activation can bind hexokinase-1 (HK-1) and result in glycolytic defects and subsequent mitochondrial dysfunction. HK-1 is the neuronal hexokinase and catalyzes the first reaction of glycolysis, converting glucose to glucose-6-phosphate (G6P), a common substrate for glycolysis, and the pentose phosphate pathway (PPP). PPP is critical in maintaining NADPH and GSH levels via G6P dehydrogenase activity. Therefore, defects in HK-1 will not only decrease cellular bioenergetics but will also cause redox imbalance due to the depletion of GSH. In brain ischemia, whether PAR-mediated inhibition of HK-1 results in bioenergetics defects and redox imbalance is not known. We used oxygen-glucose deprivation (OGD) in mouse cortical neurons to mimic brain ischemia in neuronal cultures and observed that PARP-1 activation via PAR formation alters glycolysis, mitochondrial function, and redox homeostasis in neurons. We used pharmacological inhibition of PARP and adenoviral-mediated overexpression of wild-type HK-1 (wtHK-1) and PAR-binding mutant HK-1 (pbmHK-1). Our data show that PAR inhibition or overexpression of HK-1 significantly improves glycolysis, mitochondrial function, redox homeostasis, and cell survival in mouse cortical neurons exposed to OGD. These results suggest that PAR binding and inhibition of HK-1 during OGD drives bioenergetic defects in neurons due to inhibition of glycolysis and impairment of mitochondrial function.

## 1 INTRODUCTION

PARP-1-dependent cell death plays a crucial role in brain damage after stroke.^1^ Studies have shown that PAR is a death-signaling molecule in the PARP-dependent cell death.^2^ PARP-1 utilizes NAD+ as the precursor to catalyze PAR formation on different acceptor proteins, including PARP-1 itself, via poly-ADP-ribosylation, which plays a vital role in DNA repair under normal physiological conditions. PAR is a negatively charged molecule that can also bind via electrostatic linkage to different proteins that contain PAR-binding domains.^3,4^ PAR generation is a dynamic process balanced by the activity of poly (ADP-ribose) glycohydrolase (PARG), maintaining levels of PAR in a cell under normal physiological conditions.^5,3,6^ However, when PARP is over-activated, like in brain ischemia because of large-scale DNA nicks and breaks, excessive amounts of PAR are generated.^1,7^ Under these conditions, the accumulated PAR translocates to the cytosol either as free PAR or in the form of poly-ADP-ribosylated proteins that can bind different proteins containing PAR-binding domains, resulting in the alteration of their function, structure, or localization.^7,8,1,4^ Hexokinase is a PAR-binding protein that contains a lysine/Arginine (KR) PAR-binding motif (PBM) between amino acids 422 and 441.^9,10^ PAR can bind HK and change its catalytic activity resulting in defective glycolysis and subsequent mitochondrial dysfunction.^10,9^ HK is a vital enzyme that phosphorylates glucose to glucose-6- phosphate (G6P) in the first step of glycolysis. G6P is a common substrate for glycolysis and the pentose-phosphate pathway (PPP) that maintains cellular bioenergetics via glycolysis and subsequent mitochondrial bioenergetics using glycolytically-derived pyruvate as a substrate. G6P is also required to maintain reduced nicotinamide adenine dinucleotide phosphate (NADPH) and glutathione (GSH) levels via glucose-6-phosphate dehydrogenase (G6PD) activity sustaining cellular redox homeostasis. Therefore, loss of HK activity will result in defects in cellular bioenergetics and cause redox imbalance leading to oxidative stress that contributes to pathophysiological processes accompanying PARP-1 over-activation. Our previous studies demonstrated that the binding of PAR to HK results in glycolytic and bioenergetics defects in neural cells exposed to the DNA alkylating agent, N-methyl-N’-nitro-N’-nitrosoguanidine (MNNG).^9^ PARP-1 over-activation plays a significant role in stroke-induced neural death as PARP-1 knockout (KO) mice or pharmacological inhibitors of PARP are significantly protective in brain ischemia models.^11-14^ Additionally, PARG KO mice have larger infarct lesions after middle cerebral artery occlusion, indicating that PAR likely plays a substantial role in the stroke pathophysiology.^15,2^ In mammalian cells, HK isoforms include HK-1, 2, 3, and 4 (also known as glucokinase). HK-1 is preferentially expressed in neurons and is called brain HK. Whether PAR binding to HK-1 in neurons contributes to the bioenergetic defects and redox imbalance observed during ischemia-reperfusion is not known. We show here that glycolytic defects, mitochondrial dysfunction, and oxidative stress following oxygen and glucose deprivation (OGD) in mouse cortical neurons are caused by the PAR-dependent inhibition of HK-1. Our findings suggest that PAR binding to HK-1 is a pathological event in OGD and that interfering with PAR’s interaction with HK-1 is a potential target to avert bioenergetic collapse, redox imbalance, and neuron death following ischemia.

## 2 MATERIAL AND METHODS

### 2.1 Primary mouse cortical neural cell culture

All the experiments involving the use of animals were approved by the Institutional Animal Care and Use Committee (IACUC) at the University of Alabama at Birmingham. Primary mouse cortical neurons were isolated from embryonic day 15 (E15) mouse embryos as described previously.^9,16-18^ The cortices were dissected from the E15 embryo in a dissection medium [DMEM + 20% horse serum (Thermo Fisher Scientific, Waltham, MA, USA)]. After digestion with TrypLE (Thermo Fisher Scientific) for 10 min at 37°C, the cortices were triturated using a fine pipette and straining through a 40-micron cell culture filter. The cell suspension was plated at a density of 5 × 10^5^ cells/mL on cell culture plates pre-coated with poly-L-Ornithine (Sigma-Aldrich, St. Louis, MO, USA) in Neurobasal medium [Neurobasal-A medium (Thermo Fisher Scientific) supplemented with 10 mM glucose (Sigma-Aldrich), 1 mM GlutaMax (Thermo Fisher Scientific), 1 mM sodium pyruvate (Thermo Fisher Scientific), and B-27 (Thermo Fisher Scientific). On days in vitro (DIV) 2, gilal cell growth and proliferation was inhibited by treating the cell cultures with 50μM 5-fluoro-2-deoxyuridine (Sigma-Aldrich). Mature cultures were used at DIV 11-12 for experiments and contained more than 95% neurons.

### 2.2 Oxygen-glucose deprivation (OGD) in primary cortical neural cells

OGD in primary cortical neurons was induced as previously described.^16,17,9^ The Neurobasal media was collected and saved for later use at OGD termination. The neurons were washed with PBS to remove any residual glucose. After that, OGD buffer [NaCl (116 mM), KCl (5.4 mM), MgSO_4_ (0.8 mM), NaHCO_3_ (26.2 mM), NaH_2_PO_4_ (1 mM), CaCl_2_ (1.8 mM), glycine (0.01 mM) (pH 7.4)] pre-bubbled with OGD gas (5% CO_2_, 10% H_2_, and 85% N_2_) were was added and the cultures were placed in a hypoxia chamber connected at 37°C for 90 min (Biospherix Ltd., Parish, NY, USA). The hypoxia chamber was connected to OGD gas (5% CO_2_, 10% H_2_, and 85% N_2_) to maintain the O_2_ in the chamber at less than 1% by a continuous flow of the OGD gas, measured by an oxygen sensor (Biospherix Ltd., Parish, NY, USA). OGD was terminated by oxygen-glucose resupply (OGR) achieved by adding the saved neurobasal medium back to the cultures and transferring the culture plates back to the regular incubator containing normoxic conditions (5% CO_2_ and 95% air). PARP-1 inhibitor, DPQ (30µM, Sigma-Aldrich), or vehicle (DMSO) was added to the neuronal cultures 1h before the induction of OGD.

### 2.3 Hexokinase activity assay

Hexokinase activity was assessed as described previously following Worthington protocols (Worthington, Columbus, OH, USA).^9^ Briefly, cells were lysed in cell lysis buffer [50 mM Tris (pH 7.8; Fisher Scientific), 13.3 mM MgCl_2_ (Sigma-Aldrich), 0.05% Triton X-100 (Fisher Scientific), and protease and phosphatase inhibitors (Thermo Fisher Scientific)]. For the hexokinase enzymatic reaction, 100 μL (100 μg of protein) of sample was added to 700 μL of assay buffer [50 mM Tris (pH 8.0), 13.3 mM MgCl_2_, 1 mM NAD (Sigma-Aldrich), 1 mM ATP (Sigma-Aldrich), and 1 unit/mL glucose-6-phosphate dehydrogenase (Worthington)] in a cuvette. After 5 min of equilibration at 30°C, 200 uL of 1 M glucose was added into a plate to start the enzymatic reaction. The absorbance was recorded at 1 min intervals for 10 min at 340 nm at 30°C using a spectrophotometer (DU 800, Beckman Coulter). The enzymatic activity was determined as the change in absorbance/min/mg protein.

### 2.4 Measurement of oxygen consumption ratio (OCR)

Mitochondrial OCR was measured in an XF^e^96 Extracellular Flux Analyzer (Agilent Technologies, Santa Clara, CA, USA) as previously shown.^9,17^ Briefly, neuronal cells were plated in a XF96 cell culture microplate (Agilent Technologies) at a density of 70,000 cells/well. Prior to the experiment on DIV 11, the neuronal medium was replaced with XF DMEM medium (pH 7.4) containing 10 mM glucose, 1mM L-glutamine and 1 mM Sodium Pyruvate. Neurons were incubated in this media for additional 1hr at 37^ᵒ^C in a CO_2_-free incubator. Oligomycin, carbonyl cyanide 3-chlorophenylhydrazone (CCCP), and antimycin A + rotenone (Sigma-Aldrich) were sequentially injected into each well to determine basal respiration, maximal respiration, and ATP turnover. The protocol used for the measurement was 1.5-min mix, 1-min wait, and 2-min measurement with three cycles each. The data were normalized to the protein concentration in each well and are presented as the percentage of change as compared with control.

### 2.5 Measurement of extracellular acidification rate (ECAR)

The ECAR was measured in a similar way in an XF^e^96 Extracellular Flux Analyzer ^9^, except that the neuronal medium was replaced with the XF DMEM (pH7.4) medium without glucose, glutamine, and pyruvate. After 1h incubation in a CO_2_-free incubator at 37^ᵒ^C, ECAR was measured by sequentially injecting glucose, oligomycin, and 2-deoxy-glucose (Sigma-Aldrich) following a protocol of 1.5-min mix, 30-min wait, and 2-min measurement with 3 cycles each. Basal glycolysis and glycolytic capacity were calculated and normalized to the concentration of protein in each of the corresponding microplate wells by BCA assay (Thermo Fisher Scientific).

### 2.6 Mitochondrial membrane potential

Mitochondrial membrane potential was measured by using Tetramethyl rhodamine ethyl ester (TMRE; Thermo Fisher Scientific) dye as previously shown.^17,4^ For this, the neuronal cells cultured on glass coverslips were incubated with TMRE (10nM) in neurobasal medium for 30 min in a CO_2_ incubator at 37^ᵒ^C. After incubation the cells were washed with warm PBS, and live-cell imaging was captured at 30-sec intervals on an LSM710 confocal microscope (Zeiss, Germany) equipped with a temperature-controlled stage-top incubator. After recording a baseline, 20 μM CCCP (Sigma-Aldrich) was added, and images were captured for an additional 2 min. TMRE intensity was quantified as a relative ratio ΔF/F_0_ using Zen software (Zeiss, Germany).

### 2.7 Detection of cellular oxidative stress

Cellular oxidative stress was assessed by using CellROX Green Reagent (Thermo Fisher Scientific) according to the manufacturer’s instruction. Neuronal cells grown on glass coverslips were incubated with 5 µM CellRox Green Reagent for 30 minutes at 37^ᵒ^C in the incubator.^17^ After incubation, the cells were washed with PBS. Live-cell images were captured using an LSM710 confocal microscope (Zeiss, Germany). Oxidative stress was quantified as a relative fluorescence density using Zen software (Zeiss, Germany).

### 2.8 NADP/NADPH and GSSG/GSH measurement

NADP/NADPH and GSSG/GSH were measured using NADP/NADPH colorimetric kit (BioVision, K347-100) and Glutathione Fluorometric assay kit (BioVision, K264-100), respectively, according to the manufacturer’s instruction^9^. An equal amount of protein from control and OGD samples was used for each assay to initiate the reaction. The reaction product for NADP/NADPH was measured at 450nm. Total (NADPt) and NADPH were measured. NADP was later calculated by subtracting NADPH from NADPt. For measuring GSH/GSSG, reduced glutathione (GSH) and oxidized glutathione (GSSG) was measured directly from control and OGD neurons using a fluorescence plate reader at EX/Em=340/320.

### 2.9 Adenoviral transduction in mouse primary cortical neural cells

Mouse primary cortical neural cells were transduced with control adenovirus (eGFP), wt HK-1 overexpressing adenovirus (wtHK-1) and PAR binding mutated HK-1 adenovirus (pbmHK-1); Vector Biolabs). The virus were packaged at Vector Biolabs (Malvern, PA, USA). Cells were treated with adenovirus on DIV 6 and experiments were performed on DIV 11.

### 2.10 Western blot analysis**|**

Neuronal cells were washed with ice-cold PBS and lysed in a RIPA buffer (pH 7.4, 1% Triton-X100, 0.1% SDS, 1% deoxycholate, 1mM EDTA with protease and phosphatase inhibitor). Cell lysates were separated on SDS/PAGE and proteins on the gels were transferred to nitrocellulose membranes (Bio-Rad, Hercules, CA, USA). The membranes were blocked with 5% skim milk diluted in TBST for 1 h at room temperature. After blocking, the membranes were incubated with our custom designed primary antibodies [anti-PAR; clone #19, AbD Serotec, Bio-Rad),^9^ anti-HK-1 (Cell Signaling)] overnight at 4^ᵒ^C. After incubation with the primary antibodies, the membranes were washed three times with TBST and incubated with HRP-conjugated goat anti-human IgG (Fab’)2 (Abcam) or HRP-conjugated anti-rabbit IgG (Cell Signaling) secondary antibodies for 1 h at room temperature. The membranes were visualized with SuperSignal West Pico Plus Chemiluminescent Substrate (Thermo Fisher, 34578) using a ChemiDoc MP imaging system (Bio-Rad, USA). The same blot was striped and processed to visualize β-Actin using HRP-conjugated β-Actin antibody (Abcam).

### 2.11 Immunocytochemistry

Neurons on poly-L-Ornithine coated glass coverslips were washed with ice-cold PBS and fixed in 4% (w/v) paraformaldehyde for 15 min at room temperature. Cells were thereafter permeabilized for 20 min with 0.2% (v/v) Triton X-100 in PBS containing 10% donkey serum. Cells were then blocked in blocking buffer [10% donkey (Sigma, D9663) serum in PBS] for 1 h at room temperature. Neurons were then incubated overnight at 4⁰C in primary antibodies (1: 500) (GFP, Tom20) diluted in blocking buffer. Following primary antibody incubation, cells were washed three times with PBS and incubated with the Alexa Fluor 488 and Alexa Fluor 555 conjugated secondary antibody (Thermo Fisher Scientific, 1:1000) in 1% (v/v) donkey serum in PBS for 1 h in the dark at room temperature. Afterward, cells were washed and counterstained with DAPI (300nM) for nuclear staining. After the final washes with PBS, the cells on glass coverslips were mounted onto glass slides using Immuno-Mount (Thermo Fisher Scientific). Images were captured using a Zeiss 710 Confocal Microscope (Zeiss, Germany).

### 2.12 Immunoprecipitation

Co-Immunoprecipitation of PAR and HK-1 was performed according to the method described previously.^9^ Following OGD, adenovirus transduced neuronal cells from a six-well plate (Nunc Thermo Scientific) were collected in cold PBS and centrifuged at 750g at 4 °C for 5 min to collect the neuronal cell pellet. The cell pellets were lysed in 50 µl of lysis buffer (1 mM EDTA, 1 mM EGTA, 0.5% SDS, 1% Triton-X100, with protease and phosphatase inhibitors in PBS). Thereafter, the cell lysate was re-suspended in 450µl of PBS containing protease and phosphatase inhibitors followed by incubation for 30 min at 4°C with constant agitation. The resulting cell lysates were centrifuged at 12,000 × g, 4°C for 10 min, and the supernatants were incubated with either 2 µg of anti-PAR antibody (clone #19, custom designed at AbD Serotec, Bio-Rad) or Human IgG F(ab)2 (Rockland, USA) overnight at 4°C. Then Pure ProteomeTM Kappa IgG binder magnetic beads (Millipore) were added to the antibody-lysate complex and incubated for an additional 2 h at room temperature. Next, the beads were collected and washed three times with the wash buffer (0.1% Triton-X 100 with protease and phosphatase inhibitor in PBS). The samples were eluted by heating at 100°C for 5 min in Laemmli sample loading buffer (Bio-Rad, USA), and resolved on an SDS/PAGE. Western blots were performed using anti-PAR (clone #19, custom designed at Bio-Rad, AbD Serotec GmbH), anti–HK-1 (Cell Signaling) and anti-GFP (Cell Signaling) primary antibodies, followed by HRP goat anti-human IgG (Fab’)2 (Abcam) or HRP donkey anti-rabbit IgG (Cell Signaling) secondary antibodies. The membranes were visualized with SuperSignal West Pico Plus Chemiluminescent Substrate (Thermo Fisher, 34578) according to the manufacturer’s instructions using a ChemiDoc MP Imaging System (Bio-Rad).

### 2.13 Proximity Ligation Assay (PLA)

PLA was performed with DuoLink PLA technology probes and reagents (Sigma-Aldrich, USA) according to the manufacturer’s instructions. Briefly, primary neurons were fixed with 4% paraformaldehyde (PFA) for 10 min and permeabilized with 0.2 % Triton X-100 for 20 min. After that, the cells were washed with PBS twice, incubated in the blocking solution for 30 min at 37°C, and then incubated with the primary antibodies (VDAC and HK-1) overnight at 4^ᵒ^C. Thereafter, coverslips were washed with buffer A twice (5 min each) and incubated for 60 min at 37°C with the respective PLA probes diluted in antibody diluent. After two 5 min washes with buffer A, the ligation step was performed for 30 min at 37°C with ligase diluted in ligation stock. After that, the cells were washed with buffer A twice for 2 min before incubation for an additional 100 min with amplification stock solution at 37°C. The amplification stock solution contains polymerase for the rolling circle amplification step and oligonucleotides labeled with fluorophores, which will bind to the product of the rolling circle amplification and thus allow detection. After two washes of 10 min with buffer B and one wash with PBS, the coverslips were mounted with Duolink in situ mounting medium containing DAPI. A negative control experiment was performed for every antibody in which only one antibody was incubated with the PLA probes. Images were captured using a Zeiss 710 Confocal Microscope (Zeiss, Germany).

### 2.14 Cell Death Assay

Two complementary methods were used to determine cell death in neurons.^16^ In the first method, Alamar blue (Thermo Fisher, DAL1100) reagent was used, and in the second method, Hoechst 33342 (HS) and Propidium Iodide(PI) (Thermo Fisher, USA) staining was used. For Alamar blue assay, control and OGD neurons were incubated in the neurobasal medium containing the Alamar blue reagent [10% (v/v)] for 3 h. After incubation, 100 μL of the medium from each well was transferred to a 96-well microplate. The fluorescence was measured at the excitation and emission wavelength of 540 and 595 nm, respectively, using Victor X5microplate reader (Perkin Elmer, MA, USA). For the HS and PI staining, the control and OGD neurons were incubated with 1 µM Hoechst and 5 µg/ml PI in the Neurobasal medium for 5 min. After incubation, the medium was replaced with a new medium, and images were captured using a fluorescence microscope (Axio-Observer, Carl Zeiss, Germany). HS- and PI-positive cells were counted, and cell death was calculated by diving PI-positive cells (dead cells) with the HS-positive cells (total number of cells).

### 2.15 Statistical analysis

The experimental design is illustrated in the figures (Figure 1A and 3B) and described in the figure legends and respective methodology. Neuronal cultures were randomly distributed into control and experimental groups. Statistical analyses were performed using GraphPad Prism version 9 (GraphPad, USA). Western blot quantification was performed using ChemiDoc software (Bio-Rad). Quantified data are presented as mean ± SEM. One-way analysis of variance (ANOVA) with Tukey’s post hoc test was used for multiple group comparisons.

**FIGURE 1.**
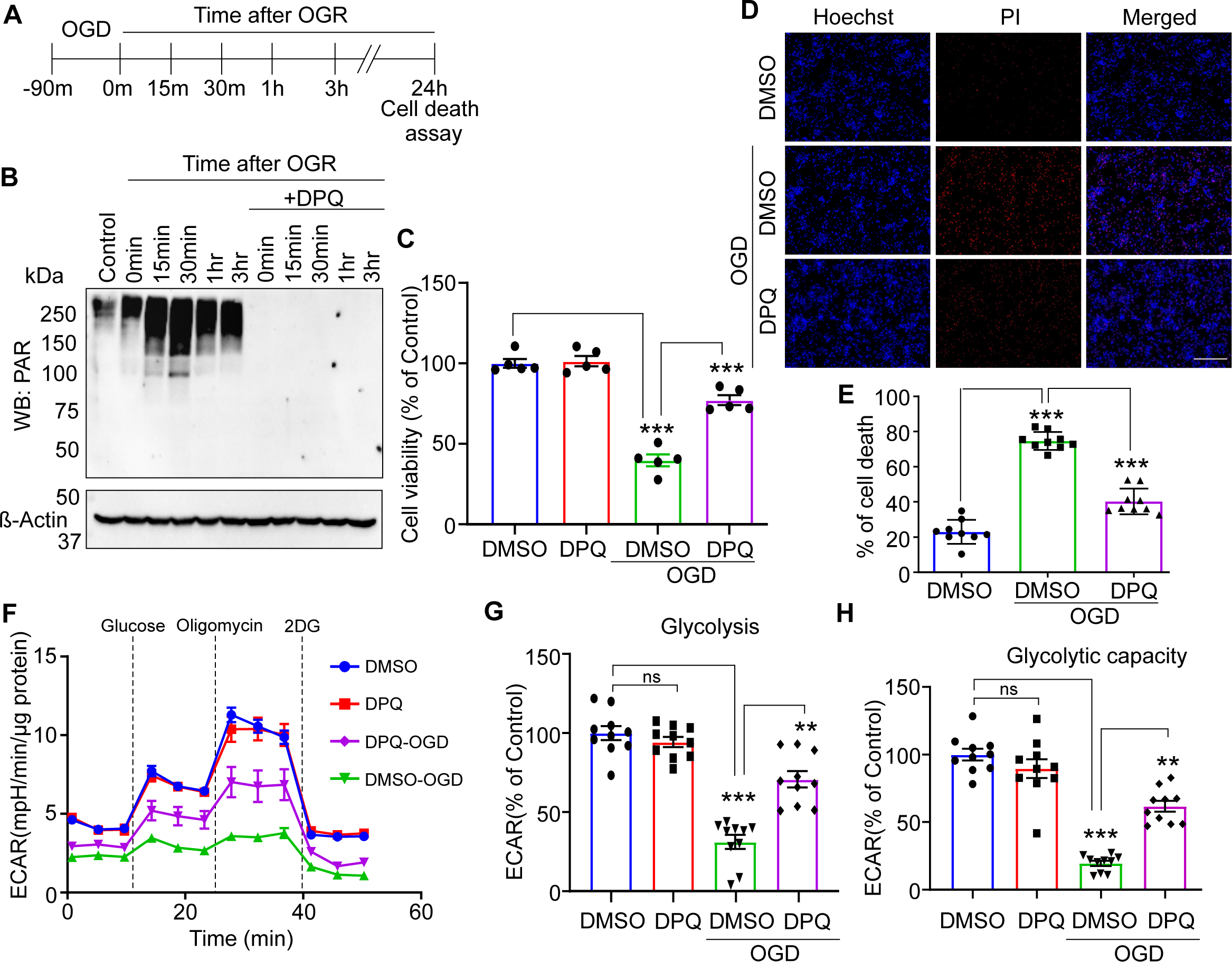
Oxygen glucose deprivation (OGD) induces neuronal death via PAR formation. (A) Schematic diagram showing the experimental design. (B) Representative western blot of PAR formation in neurons with or without DPQ after OGD at different time points of OGR. β-Actin is used as a loading control. (C) Cell viability in mouse cortical neurons under normal (control) and OGD condition in the presence or absence of DPQ (30 μM). Neurons were exposed to OGD for 90 min in the presence or absence of DPQ, and cell viability was assessed 24 h after OGR using the Alamar Blue assay (n = 5). The cell death assay were repeated three times with similar results. (D) Representative images of Hoechst 33258- and PI-labeled cells in control and OGD neurons. Scale bar: 20 μm. Primary neurons were exposed to OGD in the presence or absence of DPQ on DIV 11 and were stained with Hoechst 33258 and PI 24 h after OGR. (E) PI-positive cells were quantified from 9 cover slips collected from three independent experiments for each group and calculated as a percent of total cells. (F) ECAR analysis in control and OGD neurons with or without DPQ. (G) Glycolysis, and (H) Glycolytic capacity were calculated relative to the DMSO control (n = 10). ECAR analysis was performed three times with similar results. Data are mean ± SEM. *P < 0.05, **P < 0.01, and ***P < 0.001vs indicated groups, calculated with two-way ANOVA followed by Tukey’s post hoc test. ns, not significant.

## 3 RESULTS

### 3.1 OGD induces PARP-dependent loss of glycolysis

Brain ischemia results from the disruption of blood carrying oxygen and glucose to a particular part of the brain. Therefore, exposing neurons in culture to OGD mimics brain ischemia. We exposed mouse cortical neurons to 90 min OGD, followed by a resupply of oxygen and glucose (OGR). We collected the samples at different time points after OGR and assessed PAR formation in the presence or absence of the PARP inhibitor, 3,4-dihydro-5-[4-(1- piperidinyl)butoxy]-1(2H)-isoquinolinone (DPQ) as previously described.^9^ Our data show that OGD for 90 minutes results in PAR formation when assessed at different time points following OGR. PAR formation was detected as early as 0 min of OGR after OGD for 90 min (Figure 1A,B). DPQ- completely inhibited the PAR formation in these neurons (Figure 1B). Assessment of cell death 24 h after the termination of OGD further demonstrated that cell death following OGD is PARP-dependent as the PARP inhibitor, DPQ, significantly protected against cell death (Figure 1C,D,E). These data are consistent with prior observations that cell death in OGD is PARP-dependent.^19,20^ Previous studies showed that PAR inhibits glycolysis and cellular bioenergetics in MNNG-treated mouse cortical neurons.^9^ In a similar manner, glycolysis was assessed as the extracellular acidification rate (ECAR) in the presence of glucose (10 mM) in OGD-exposed mouse neuronal cultures in the presence or absence of DPQ using an XFe96 Seahorse flux analyzer (Agilent Technology, USA). Similar to MNNG-treated neurons^9^, OGD results in loss of glycolysis that was significantly prevented by inhibition of PARP with DPQ (Figure 1F,G,H). DPQ prevented the reduction in both basal and maximal glycolysis in OGD-exposed mouse cortical neurons (Fig. 1F,G,H).

### 3.2 PARP inhibition preserves mitochondrial function in OGD-exposed neurons

Glycolysis-derived pyruvate is a principal substrate for the mitochondrial tricarboxylic acid (TCA) cycle and mitochondrial bioenergetics in neural cells. To determine whether OGD-mediated defects in glycolysis also result in mitochondrial dysfunction, we assessed mitochondrial function as oxygen consumption rate (OCR) in mouse neuronal cultures exposed to OGD. OCR was measured using an XFe96 Seahorse Flux analyzer in mouse cortical neurons after 1 h of OGD termination in seahorse media containing 10 mM glucose, 1 mM glutamine, and 1 mM pyruvate. Oligomycin (1 µM), CCCP (1µM), and rotenone/antimycin A (1µM each) were sequentially added to assess basal respiration, maximal respiration, and ATP turnover. OGD significantly decreased mitochondrial function in mouse cortical neurons, which was significantly prevented by inhibition of PARP with DPQ (Figure 2A). Basal OCR, maximal OCR, and ATP turnover were decreased in OGD-exposed neurons, which was significantly prevented by DPQ (Figure 2B,C,D). To assess whether the OGD-mediated inhibition of glycolysis (see Figure 1) was responsible for the decrease in mitochondrial OCR in OGD-exposed neurons, the glycolytic process was by passed via supplementation with pyruvate (10 mM) directly to the neuronal cultures at the time of OGD termination. Supplementation with 10 mM pyruvate at the time of OGD termination significantly prevented the mitochondrial OCR deficits in OGD-exposed mouse primary neurons (Figure 2E,F,G,H). These data taken together suggest that PAR-induced inhibition of glycolysis caused cellular bioenergetic defects in neurons after OGD.

**FIGURE 2.**
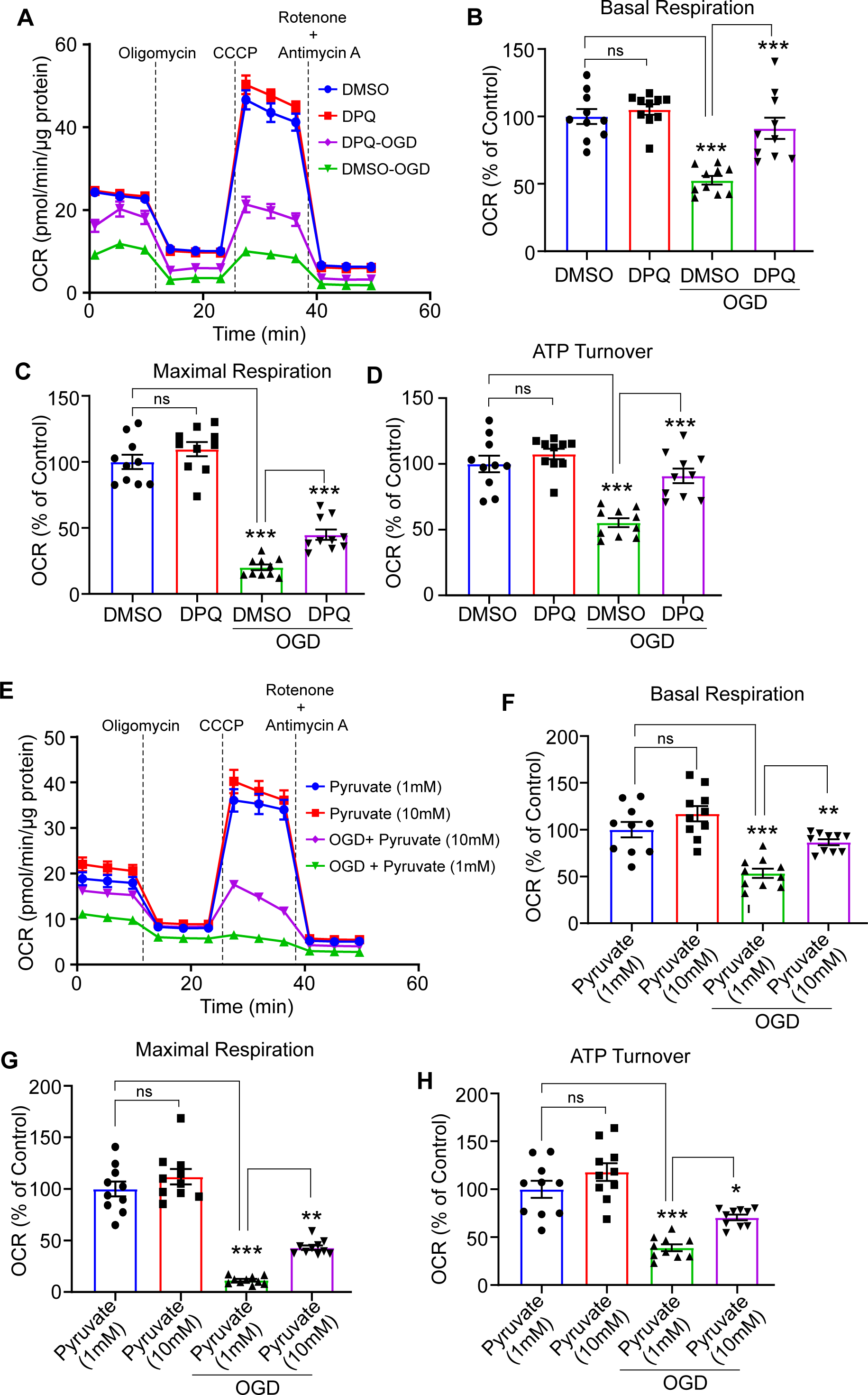
Inhibition of PAR improves mitochondrial function. (A) Representative OCR profile of mouse cortical neurons after OGD with or without DPQ treatment (n = 10). (B) Basal respiration (C) Maximal respiration and (D) ATP turnover. All these parameters were calculated relative to the DMSO control (n = 10). OCR experiments were repeated three times with similar results. (E) Representative OCR in control and OGD neuron in presence of low (1mM) or high (10mM) pyruvate (n = 10). (F) Basal respiration (G) Maximal respiration and (H) ATP turnover. All these parameters were calculated relative to the DMSO control (n = 10). OCR experiments were repeated two times with similar results. Data are mean ± SEM. ***P < 0.001vs indicated groups, calculated with two-way ANOVA followed by Tukey’s post hoc test. ns, not significant.

### 3.3 PAR binds HK-1 and decreases HK-1 activity in OGD

Mouse cortical neurons were exposed to OGD, and 1 h after OGD terminated, samples were collected to determine HK activity. OGD led to a significant decrease in HK activity that was prevented by the PARP inhibitor DPQ (Figure 3A). No change in HK-1 levels was observed (Figure 3B), indicating that the changes in HK activity are due to changes in the catalytic activity of HK-1 in mouse cortical neurons. Immunoprecipitation of PAR indicates that HK-1 interacts with PAR in OGD-exposed neurons (Figure 3C). PAR binds to a number of PAR binding domains, where PAR binding can, in part, inhibit or activate a protein.^6^ HK-1 and HK-2 contain a KR motif PAR binding domain,^21,22^ amino acids ^422^-QYS<u>RR</u>FH<u>K</u>TL<u>RR</u>LVPDSDVR-^441^.^9^ Individual HK-1 peptides containing the KR motif PAR binding domain with the replacement of four arginine (R) and one lysine (K) sequentially were synthesized and spotted on a modified nitrocellulose membrane array using Intavis Spot technology to characterize the PAR-binding domain. The membrane was hydrated in methanol and equilibrated in Tris-buffered saline, followed by incubation with free ^32^P-labeled PAR purified on DHBB resin. The membrane was extensively washed in TBS, dried, autoradiographed on a Fuji imaging Film, and assessed with a Typhoon imager. PAR was observed to bind to the HK-1 peptides, and the PAR binding became weaker as the individual R or the K amino acids were replaced by alanine. Complete loss of PAR binding was observed when all four R and the K were changed to alanine (A) (Figure 3D). A PAR Binding mutant HK1 (pbmHK-1) was generated by replacing the four R and one K to As, followed by packaging into adenoviral particles (Vector Biolabs, USA). Mouse cortical neurons were transduced with eGFP-tagged wild-type HK-1 (wtHK-1), PAR binding mutated HK-1 (pbmHK-1), or eGFP (control) adenoviruses on DIV 6, and expression was assessed on DIV 11. eGFP-tagged pbmHK-1 or wtHK-1 displayed efficient transduction and expression in mouse cortical neurons on DIV 11 (Figure 4A,B). Following exposure to OGD for 90 min on DIV 11, the neuronal samples were subjected to PAR immunoprecipitation with an anti-PAR antibody.^9^ PAR co-immunoprecipitates wtHK-1 or endogenous HK-1, whereas PAR is unable to co-immunoprecipitate pbmHK-1 (Figure 4C). These data confirmed that HK-1 contains a PAR-binding domain,^9^ and further identified that pbmHK-1 is devoid of PAR binding.

**FIGURE 3.**
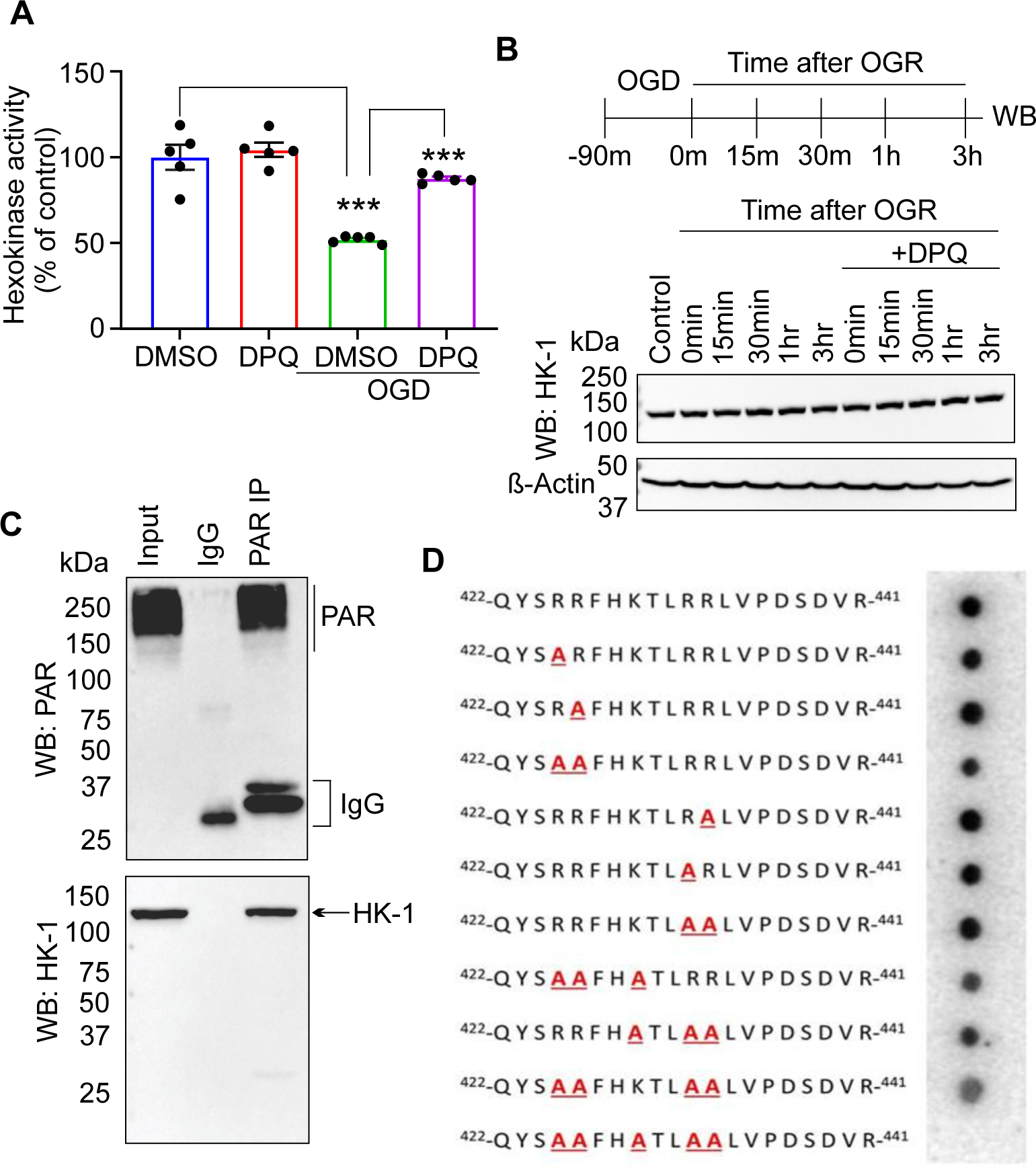
PAR inhibits HK activity in neurons. (A) HK activity assay in mouse cortical neurons with or without DPQ (30 μM) under normal and OGD condition (n = 6). Data are expressed as mean ± SEM. **P < 0.01 and ***P < 0.001vs indicated groups, calculated with two-way ANOVA followed by Tukey’s post hoc test. (B)Top: Schematic diagram showing the experimental design. Representative western blot showing levels of HK-1 in neurons with or without DPQ after OGD at different time points of OGR. β-Actin is used as a loading control. (C) Western blot data showing co-immunoprecipitation of endogenous HK-1 with PAR. PAR is observed in inputs and PAR-pull down samples. PAR IP was performed using a custom-made PAR monoclonal antibody (Materials and Method) from mouse cortical neurons after OGD. HK-1 blot developed with HK-1 antibody on PAR blot. (D) PAR dot blot performed with peptides containing PAR- binding motifs or sequentially deleted PAR-binding residues of HK-1 and recombinant PAR.

**FIGURE 4.**
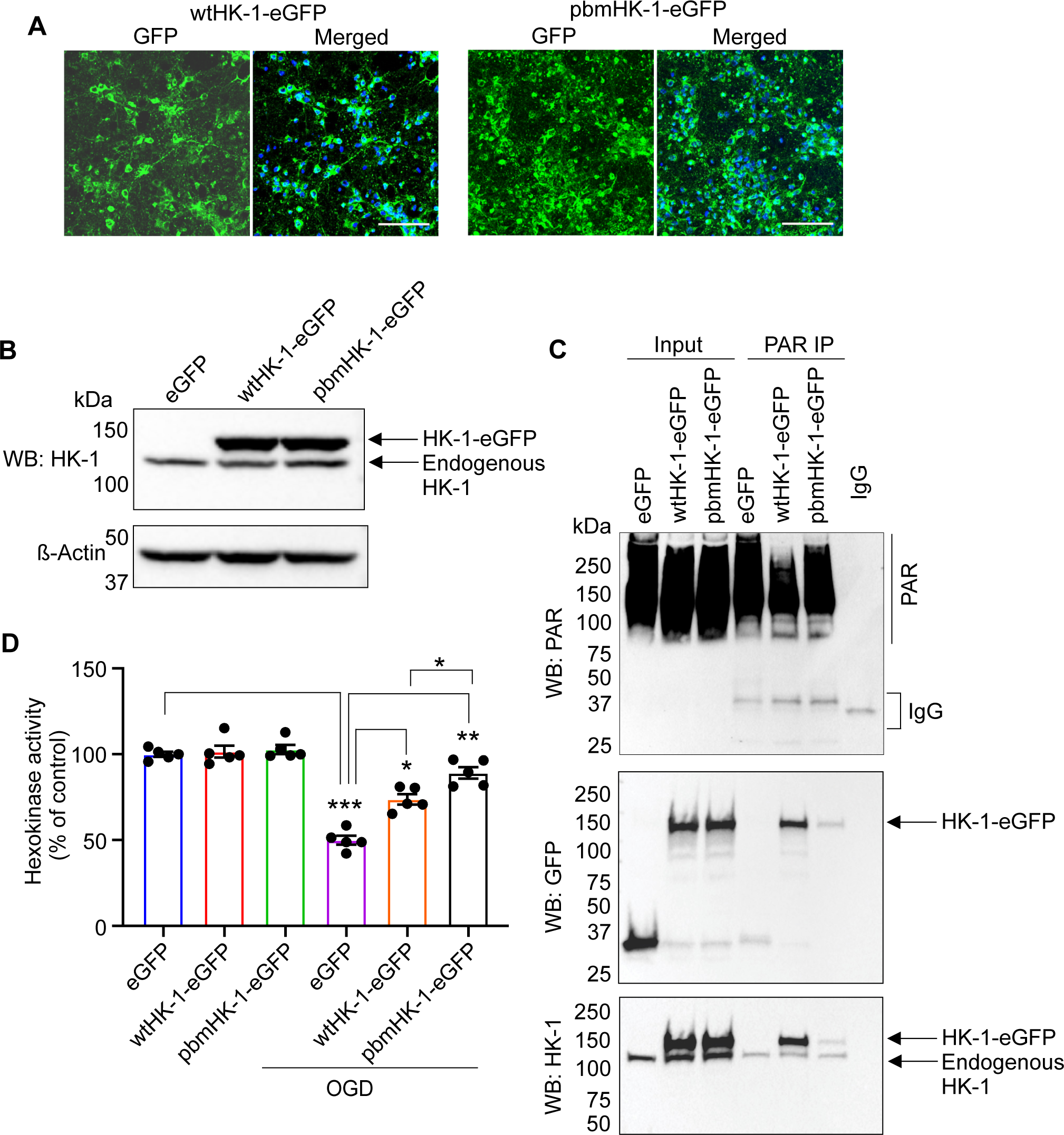
PAR binding sited mutated HK-1(pbmHK-1) does not bind with PAR and is more active than WT HK-1 in neuron after OGD. (A) Confocal images showing transduction efficiency of adenovirus carrying wtHK-1 and PAR-binding mutated HK-1 (pbmHK-1) in primary neurons. Scale bar: 20 μM. (B) Western blot of wtHK-1 and PAR-binding site mutated HK-1 (pbmHK-1) expression in neurons transduced with respective adeno viruses. Upper bands on the blots represent adeno virus mediated exogenous HK-1 and lower bands represent endogenous HK-1. Neurons were transduced with respective adenoviruses on DIV 6 and analyzed on DIV 11. (C) Top: Western blot showing PAR in inputs and IP samples obtained from neurons transduced with adenovirus of eGFP (Control), wtHK1-eGFP (wt HK-1) and pbmH1-eGFP (PAR binding mutated HK-1). A custom-made PAR monoclonal antibody was used to pull down PAR from OGD neurons. Middle: Western blot developed on the same blot using antibody against GFP, showing interaction of PAR with wtHK-1 - but not with PAR biding site mutated HK-1 (pbmHK-1- eGFP). Bottom: Western blot with HK-1 antibody on inputs and PAR IP samples. (D) Hexokinase activity assay in control and OGD neurons transduced with eGFP, wtHK1-eGFP and pbmHK1-eGFP. Data are mean ± SEM (n=5). *P < 0.05, **P < 0.01 and ***P < 0.001vs indicated groups, calculated with two-way ANOVA followed by Tukey’s post hoc test.

In subsets of neuronal cultures, HK activity was measured following transduction of the cultures with wtHK-1, pbmHK-1, or eGFP control adenoviruses. HK activity was significantly reduced in OGD-exposed GFP control adenovirus transduced neurons (Figure 4D), while overexpression of both wtHK-1 or pbmHK-1 via adenoviral transduction significantly prevented the reduction in HK activity (Figure 4D). These results taken together indicate that PAR binds to HK-1 via a KR motif PAR-binding domain where it inhibits HK-1 activity, and PAR dependent inhibition of HK-1 can be prevented by overexpression of either WTHK-1 or pbmHK-1.

### 3.4 HK-1 overexpression recovers glycolysis in OGD-exposed neurons

Glycolytic flux after exposure to OGD was assessed after overexpression of wtHK-1 or pbmHK- 1 via adenoviral transduction in mouse cortical neurons. ECAR analysis via an XFe96 flux analyzer was utilized and demonstrated that both wtHK-1 and pbmHK1 significantly prevented the defects in basal glycolysis and glycolytic capacity as compared to adeno GFP transduction in neurons exposed to OGD (Figure 5A,B,C). The effect on glycolysis in neurons transduced with pbmHK-1 was significantly higher than in neurons transduced with wtHK-1. The significant recovery by wtHK-1 is likely because overexpression compensates for the PAR-mediated inhibition of endogenous HK-1. Taken together, these results suggest that PAR-mediated inhibition of HK-1 is a primary cause of glycolytic defects in OGD.

**FIGURE 5.**
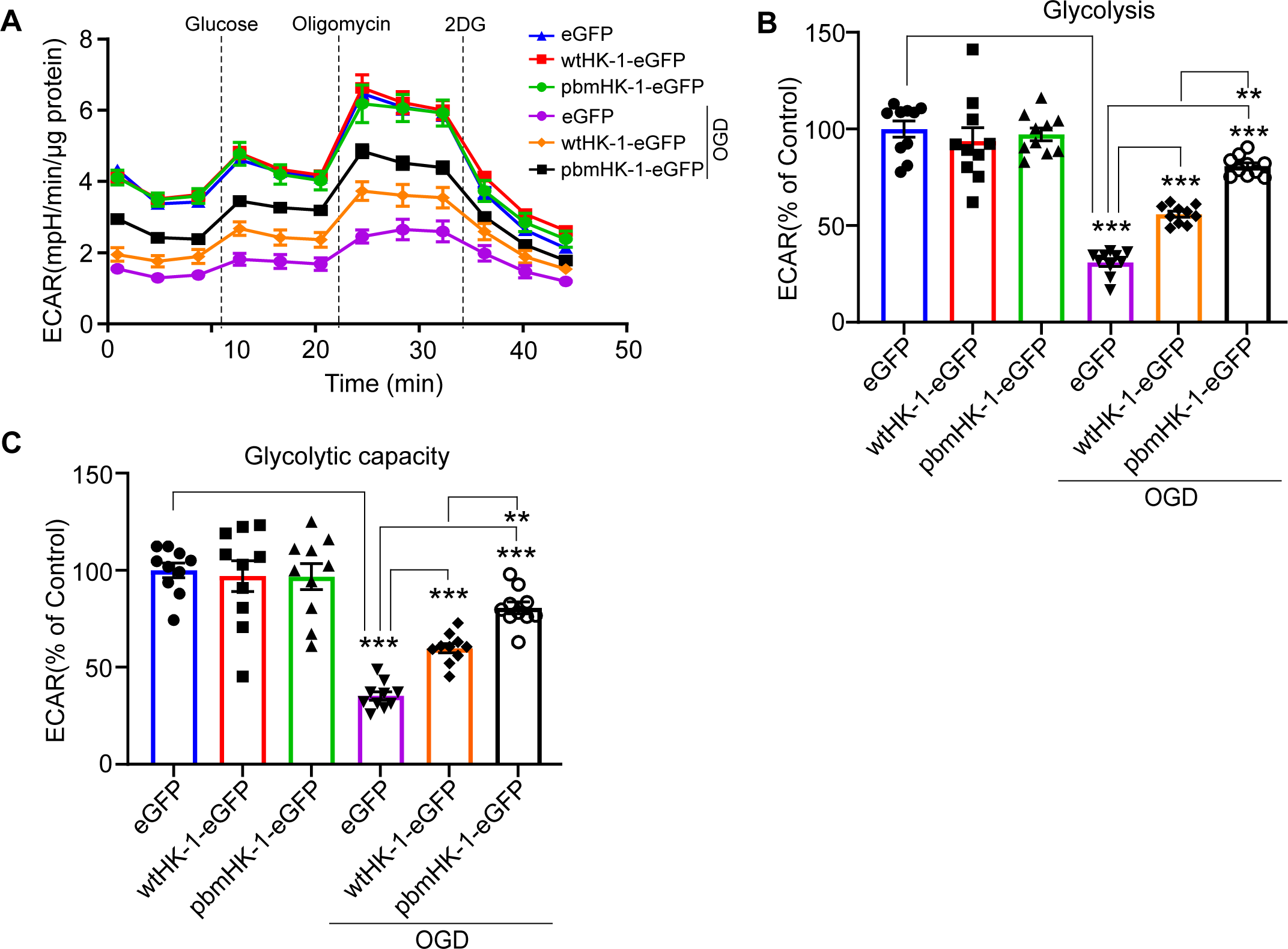
Assessment of the effect of wtHK-1 and PAR binding site mutated HK-1 (pbmHK-1) on ECAR in OGD neurons. (A) ECAR analysis in control and OGD neurons following transduction with Adenovirus of eGFP, wtHK-1-eGFP or pbmHK-1-eGFP virus. (B) Glycolysis and (C) Glycolytic capacity in control and OGD neurons extracted from figure (A) (n = 10). ECAR experiments were repeated three times with similar results. Data are expressed as mean ± SEM. **P < 0.01 and ***P < 0.001vs indicated groups, calculated with two-way ANOVA followed by Tukey’s post hoc test.

### 3.5 HK-1 overexpression improves mitochondrial function in OGD-exposed neurons

Mitochondrial function after OGD was assessed in cortical neurons transduced with wtHK-1, pbmHK-1 or GFP control adenoviruses via OCR using an XF96e flux analyzer (Figure 6A). Mitochondrial basal OCR, maximal OCR, and ATP turnover were significantly higher in neurons transduced with pbmHK1 compared to neurons transduced with GFP control virus following OGD (Figure 6B,C,D). Overexpression of wtHK-1 also significantly improved mitochondrial OCR as compared to control GFP transduced neurons following OGD (Figure 6A). The effect with wtHK-1 was significantly lower when compared to pbmHK-1 expressing neurons (Figure 6A). These data indicate that recovery of hexokinase activity via overexpression can recover mitochondrial function in neurons following OGD and indicate that PAR-mediated inhibition of HK-1 may be one of the primary processes leading to a decrease in mitochondrial function in OGD.

**FIGURE 6.**
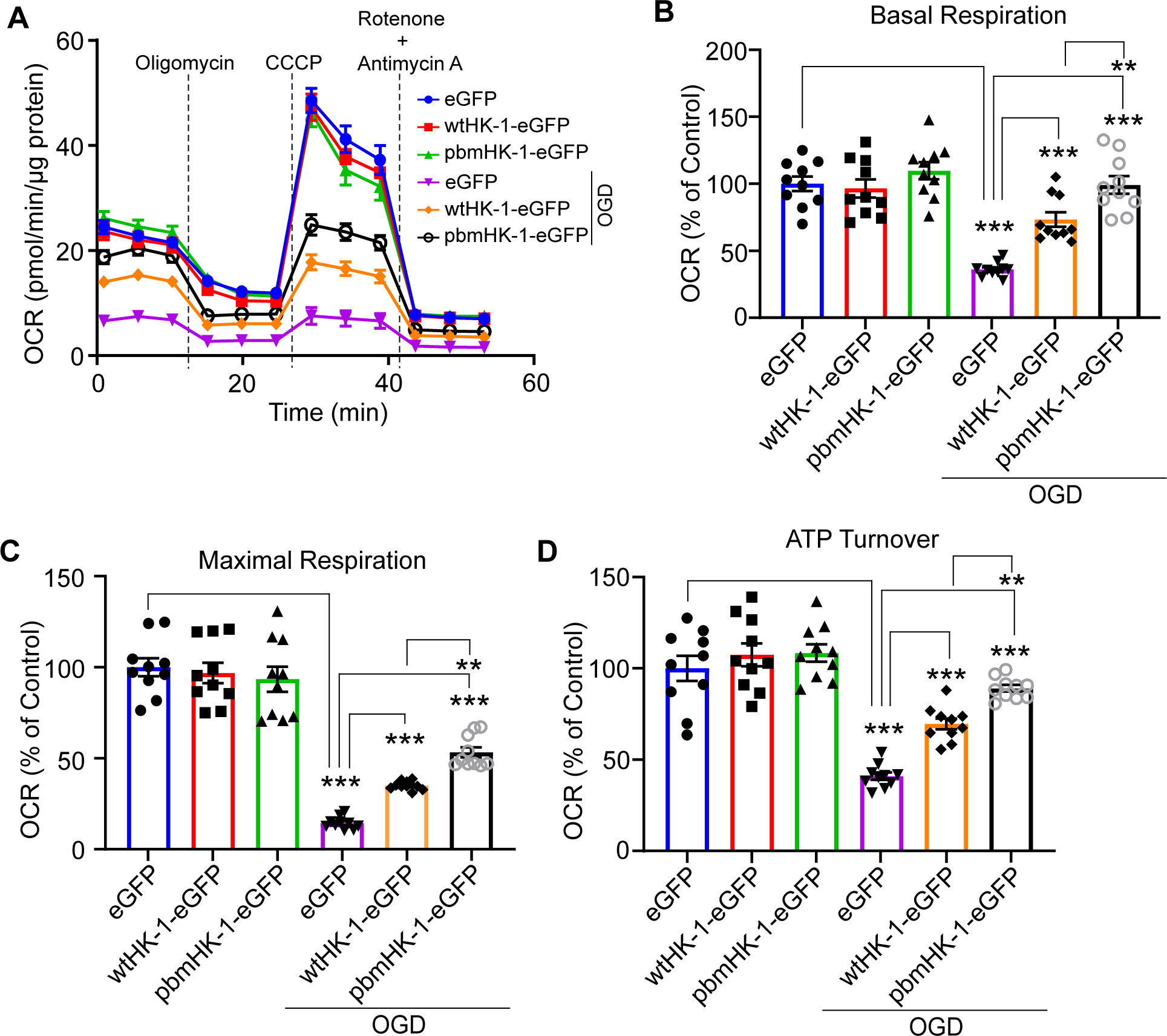
Effect of wtHK-1 and PAR binding site mutated HK-1 (pbmHK-1) on mitochondrial function in OGD neurons. (A) OCR analysis in control and OGD neurons following transduction with GFP, wtHK-1-eGFP or pbmHK-1-eGFP virus. (B) Basal respiration, (C) Maximal respiration and, (D) ATP turnover capacity in control and OGD neurons extracted from figure (A) (n = 10). Experiments were repeated three times with similar results. Data are mean ± SEM. **P < 0.01 and ***P < 0.001vs indicated groups, calculated with two-way ANOVA followed by Tukey’s post hoc test.

### 3.6 OGD-induced PARP activation causes oxidative stress in cortical neurons

Reactive oxygen species (ROS) production after OGD was monitored in mouse cortical neurons transduced with wtHK-1, pbmHK-1, or GFP control adenoviruses or in the presence/absence of the PARP inhibitor, DPQ. OGD caused ROS generation, which was significantly decreased by DPQ (Figure 7A,B). Likewise, overexpression of wtHK-1 and pbmHK-1 via adenoviral transduction significantly reduced ROS levels in neurons following OGD. Although the levels of ROS were significantly lower in both wtHK-1 and pbmHK-1 as compared to GFP control-expressing neurons, the effects on ROS reduction were greater in pbmHK-1-expressing neurons as compared to neurons transduced with wtHK-1 (Figure 7C,D). Protection against ROS production by overexpression of both wtHK-1 and pbmHK-1 suggests that PARP-dependent ROS production in OGD is mediated through inhibition of HK-1 by PAR; and both wtHK-1 and pbmHK-1 are capable of compensating for the loss of endogenous HK activity.

**FIGURE 7.**
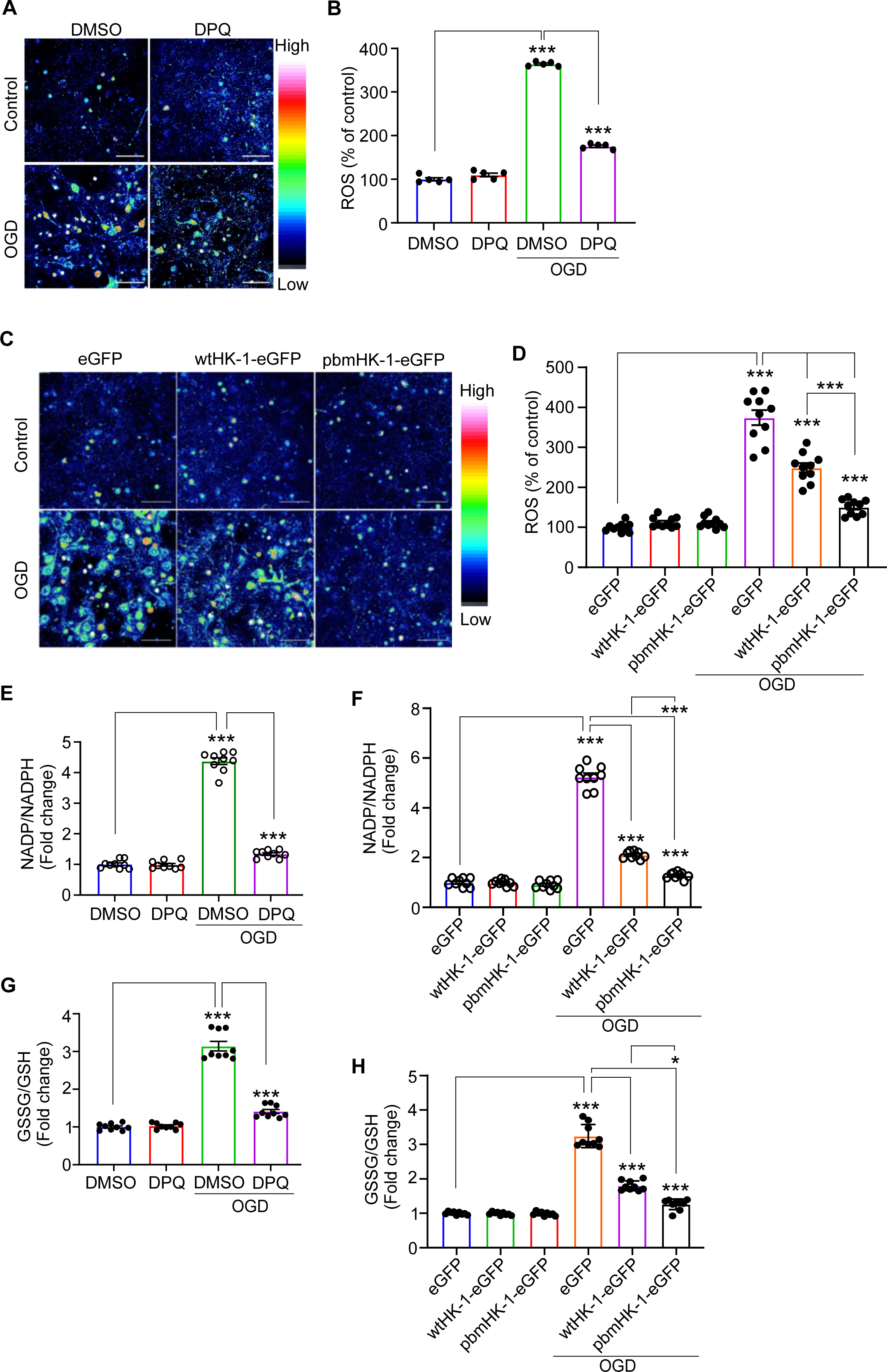
Assessment of the effect of PAR inhibition on reactive oxygen species and NADP/NADPH and GSSG/GSH in neurons after OGD. (A) Representative confocal images of CellRox fluorescence in control and OGD neurons with or without DPQ. Scale bar: 20 μm. (B) Quantification of average CellRox fluorescence intensity (ROS) collected from five fields from five experiments. (C) Confocal images of ROS generation in control and OGD neurons transduced with eGFP, wtHK-1-eGFP, or pbmHK-1-eGFP virus by CellRox staining. Scale bar: 20 μm. (D) Quantification of ROS determined as average CellRox fluorescence intensity from obtained from ten fields from five experiments. Quantification of NADP/NADPH ratio (oxidized vs reduced form) in control and OGD neurons in presence or absence of DPQ (E) and in presence of eGFP, wtHK-1-eGFP or pbmHK-1-eGFP virus (F). Data represent fold change of control and n = 9 (collected from three different experiments). Quantification of GSSG/GSH ratio (oxidized vs reduced form of glutathione) in control and OGD neurons in presence of or absence of DPQ (G) and in presence of eGFP, wtHK-1-eGFP or pbmHK-1-eGFP virus (H). Data are presented as fold change of control and n = 9 (collected from three different experiments). Data are mean ± SEM. **P < 0.01 and ***P < 0.001vs indicated groups, calculated with two-way ANOVA followed by Tukey’s post hoc test.

HK-1 in neurons catalyzes the first reaction in glycolysis, leading to the formation of G6P from glucose.^23^ G6P is a common substrate for glycolysis and the PPP.^24^ In PPP, G6P is catalyzed by G6PD into 6-phosphogluconolactone. In the process, G6PD uses nicotinamide adenine dinucleotide phosphate (NADP) as a co-factor and converts it into NADPH, which is required to convert oxidized glutathione (GGSG) back into reduced GSH by glutathione reductase.^25,26^ Therefore, defects in HK-1 in neurons will result in loss of G6P and ultimately decreased NADP and GSH. Accordingly, NADPH and GSH was measured in mouse cortical neurons following OGD, respectively. OGD caused a decrease in the levels of NADH and GSH in mouse cortical neurons, which was significantly prevented by the PARP inhibitor, DPQ. (Figure 7E,G). In a similar manner, overexpression of pbmHK-1 or wtHK-1 recovers both NADPH and GSH in mouse cortical neurons exposed to OGD (Figure 7F,H). These data indicate that PAR-mediated inhibition of HK-1 in mouse cortical neurons not only inhibits glycolysis (see Figure 5) but also results in decreased PPP flux that facilitates oxidative stress via depletion of NADPH and GSH levels.

### 3.7 PAR leads to the dissociation of HK-1 from mitochondria

HK-1 in neurons is a mitochondrial-associated protein that binds VDAC on the outer mitochondrial membrane.^27-29^ Proximity Ligation Assay (PLA) was used to assess the interaction between VDAC and HK-1 using Duolink PLA Kit (Sigma). OGD significantly reduces the interaction between VDAC and HK-1, as assessed by PLA (Figure 8A,B). Inhibition of PARP by DPQ significantly prevents the VDAC and HK-1 disassociation induced by OGD (Figure 8A,B). The association of HK-1 with mitochondria is crucial for cellular bioenergetics and vital for protecting mitochondrial membranes against the permeability transition.^30^ One of the consequences of mitochondrial permeability transition is the loss of mitochondrial membrane potential (Δψ_m_). Δψ_m_ was assessed using TMRE live-cell imaging. OGD caused a significant loss of Δψ_m_, which was prevented by the PARP inhibitor, DPQ (Figure 8C,D,E). To assess whether the loss of Δψ_m_ is due to PAR-mediated alteration/dissociation of HK-1, mouse cortical neurons were transduced with control (GFP), wtHK-1 or pbmHK-1 adenoviruses and Δψ_m_ via TMRE live cell imaging was assessed after OGD treatment. The Δψ_m_ was preserved in neurons with overexpression of wtHK-1 or pbmHk-1 (Figure 8F,G,H). Although the overexpression of both wtHK-1 and pbmHK-1 caused significant protection against OGD-mediated loss of Δψ_m_, the effect was more evident in pbmHK-1 compared to wtHK-1 expressing neurons (Figure 8F,G,H). These data indicate that PAR-binding to HK-1 in mouse cortical neurons also dissociates HK-1 from mitochondria, making mitochondria more susceptible to permeability transition.

**FIGURE 8.**
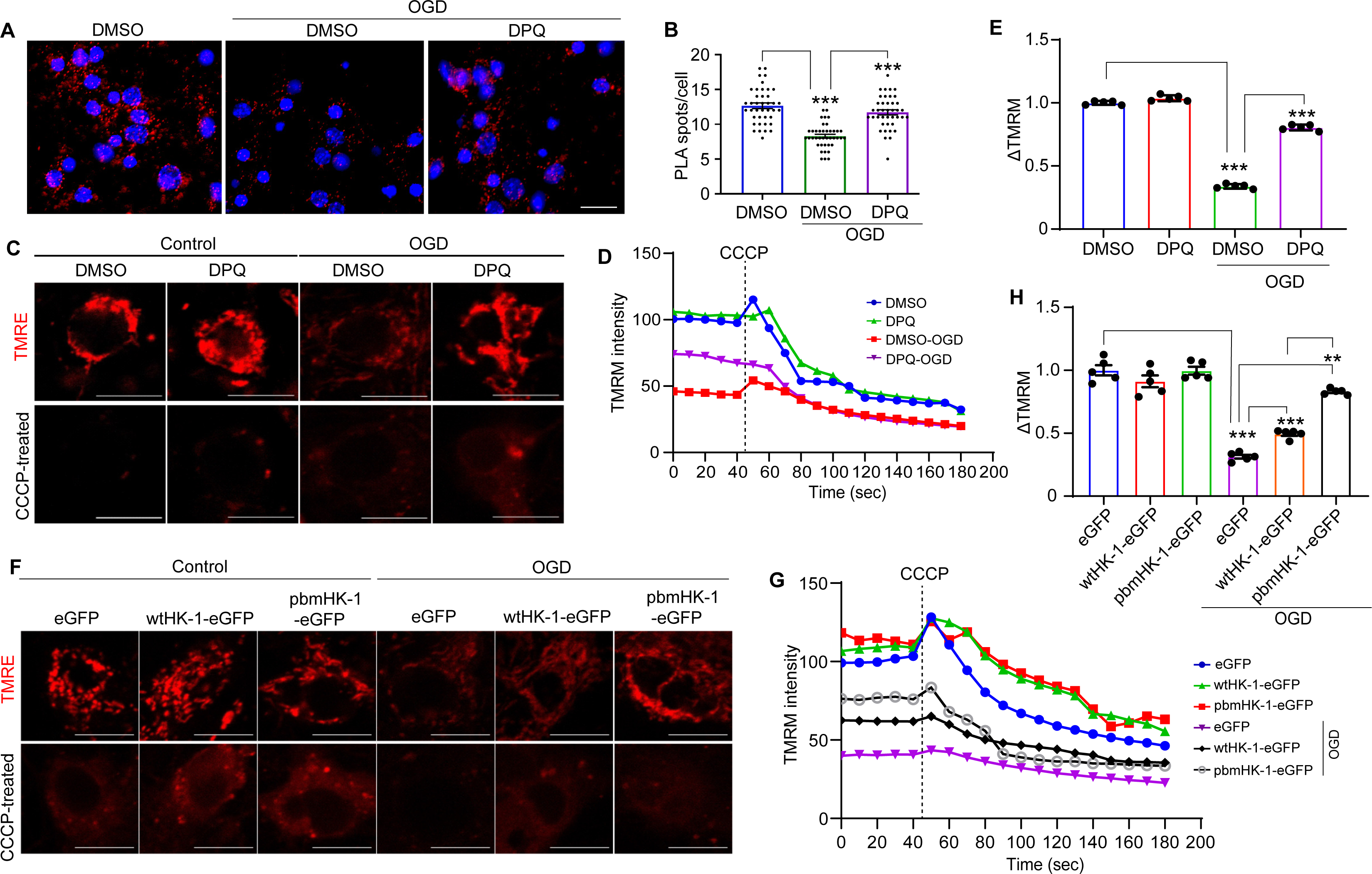
Inhibition of PAR improves mitochondrial membrane potential. (A) Representative confocal images of Proximity Ligation Assay (PLA) with anti HK-1 and anti-VDAC antibodies in control and OGD neurons with or without DPQ. Red fluorescence spots indicate association of HK-1 and VDAC. Scale bar = 20 μm. (B) Quantification of PLA spots obtained from figure (A). PLA spots were calculated from five separate experiments. (C) Representative confocal images showing TMRE intensity in mouse cortical neurons under normal (control) and OGD condition in presence or absence of DPQ (30 μM). Scale bar = 20 μm. (D) Time-lapse quantification of TMRE fluorescence intensity before and after CCCP addition. (E) Bar graph showing quantification of TMRE intensity change (ΔTMRE) before and after CCCP addition in neurons shown in figure (A) (n = 3). (F) Representative confocal images showing TMRE intensity in control and OGD neurons transduced with eGFP, wtHK-1-eGFP or pbmHK-1-eGFP adenovirus. Scale bar = 20 μm. (G) Time-lapse quantification of TMRE fluorescence intensity before and after CCCP addition obtained from figure (F). (H) Quantification of TMRE intensity change (ΔTMRE) before and after CCCP addition in neurons shown in figure (F) (n = 3). Data are mean ± SEM. **P < 0.01 and ***P < 0.001vs indicated groups, calculated with two-way ANOVA followed by Tukey’s post hoc test.

### 3.8 Overexpression of HK-1 is protective against OGD-mediated cell death

Since overexpression of wtHK-1 or pbmHK-1 protects against loss of glycolysis, mitochondrial OCR, reduces ROS and preserves Δψ_m_, we assessed whether wtHK-1 or pbmHK-1 could protect against OGD-induced cell death in mouse cortical neurons. Mouse cortical neurons were transduced wtHK-1, pbmHK-1, or control adenoviruses on DIV 6 and exposed to 90 min OGD on DIV 11. Cell death/survival was assessed via two different but complementary methods, Alamar blue assay and quantification of Propidium Iodide/Hoechst staining 24 h after OGD terminated. Overexpression of both wtHK-1 and pbmHK-1 protects neurons against OGD-mediated cell death (Figure 9A,B,C). The protection is more robust in neurons with pbmHK-1 overexpression.

**FIGURE 9.**
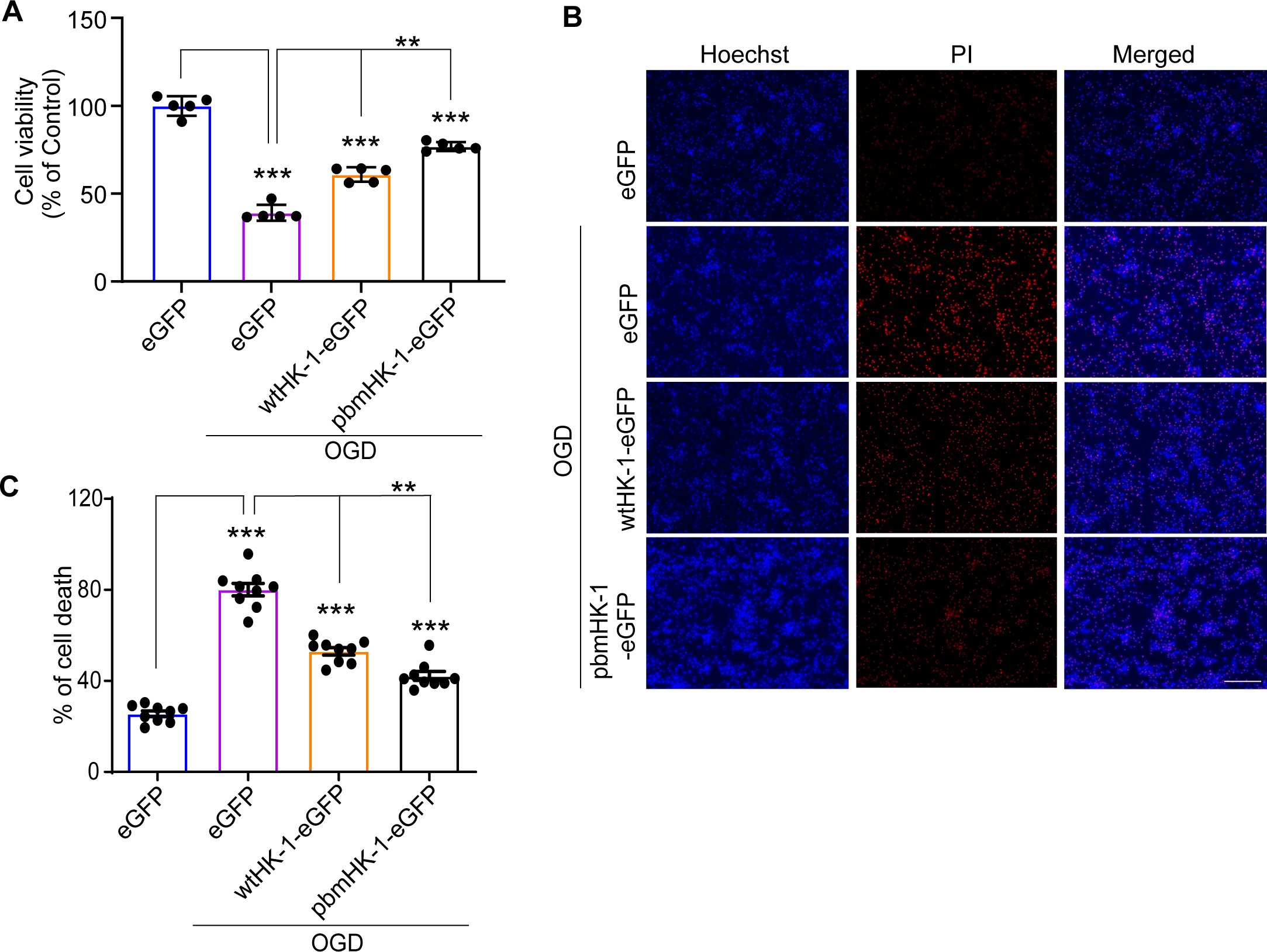
Effect of wt HK-1 and PAR binding site mutated HK-1 (pbmHK-1) on neuronal survival. (A) Cell viability in control and OGD neurons transduced with eGFP, wtHK-1-eGFP or pbmHK-1-eGFP adenovirus. Neurons were transduced with virus on DIV 6. Cell viability was measured on DIV 11 using the Alamar Blue assay (n = 5). The assay was repeated three time similar results. (B) Representative images of Hoechst 33258- and PI-labeled cells in control and OGD neurons with adenoviruses mentioned in figure (A). Scale bar: 20 μm. (C) PI-positive cells were quantified from 9 cover slips collected from three independent from experiments for each group and calculated as a percent of total cells. Data are mean ± SEM. **P < 0.01 and ***P < 0.001vs indicated groups, calculated with two-way ANOVA followed by Tukey’s post hoc test.

## 4 DISCUSSION

This study reports that PAR-mediated alteration on HK-1 results in glycolytic and mitochondrial defects in neurons exposed to OGD, a model that mimics brain ischemia in neuronal cultures. OGD resulted in a dramatic decrease in neuronal glycolysis (basal glycolysis and glycolytic capacity), which was significantly reversed by PARP inhibition, suggesting that PARP activation is a major pathological pathway leading to glycolytic defects in brain ischemia. In addition, OGD caused a PARP-1 dependent decrease in mitochondrial function as assessed by OCR (basal OCR, maximal OCR, and ATP turnover). The reduction in glycolytic flux was likely responsible for the decrease in mitochondrial OCR as in neurons, glycolytically derived pyruvate forms the principal substrate for the mitochondrial TCA function.^31,32^ Our previous studies showed that reduction in NAD+ depletion following PARP activation is not the primary cause of glycolytic defects in PAR activation, but it is PAR-dependent alteration in glycolytic enzymes, notably HK- 1 in the case of neurons.^9^ Our results agree with our previously published studies that glycolytic defects independent of NAD+ depletion are the primary cause of cellular bioenergetic failure following the PARP activation.^10^ Since Pyruvate significantly improved the neuronal mitochondrial OCR (basal OCR and maximal OCR), it is likely that the initial glycolytic defects are in part mediated by PAR-dependent defects in glycolysis and glycolytic defects precede the mitochondrial dysfunction in OGD exposed neurons. Our results do not exclude the role of NAD+-depletion in mediating the bioenergetics defects in OGD or stroke, but the NAD+- dependent pathways seem to be secondary to the initial PAR-mediated alterations in glycolysis. HK-1 is the neuronal HK,^33^ and our results confirmed that PARP is activated in OGD and PAR binds HK-1 and likely inhibits its activity as reported previously.^9,10^ Indeed, our data show that hexokinase activity is decreased in neurons exposed to OGD, which is prevented by pharmacological inhibition of PARP, revealing that the inhibition of HK activity is likely due to PAR-dependent inhibition of HK-1. Our data also show that the decrease in activity is most likely via inhibition of HK-1 catalytic activity and not due to the decrease in the levels of HK-1 in neurons.

HK-1 and HK-2 contain a KR motif PAR binding domain^22,21^ that provides a net positive charge making the electrostatic interaction with PAR (negatively charged molecule) viable.^9^ Our peptide dot blot data revealed that the PAR-binding capacity of HK-1 could be abolished only if all the lysine and arginine amino acids in the PAR-binding motif were mutated to alanine. Expression of pbmHK-1 (PAR-binding mutant HK-1) in mouse cortical neurons prevented the inhibition of HK activity after OGD. Overexpression of wtHK-1 also prevented the inhibition of HK activity, improving glycolytic flux and mitochondrial function, suggesting that overexpression of HK-1 was capable of compensating for the loss of endogenous HK due to PAR inhibition in OGD exposed neurons. The enhanced HK activity engendered by pbmHK-1 versus wtHK-1 is consistent with the notion that HK binding by PAR inhibits HK activity. These data are consistent with the idea that PAR-mediated inhibition of HK-1 mediates the bioenergetics dysfunction in brain ischemia.

HK-1 catalyzes the formation of G6P from glucose in the first step of glycolysis.^34^ At this step, glucose metabolism branches into glycolysis and PPP making G6P a common substrate for both these pathways. Defects in hexokinase activity will affect cellular glycolysis and PPP.^35,36^ The first step in PPP is vitally required to maintain GSH levels in the cells via NADPH.^37^ A decrease in hexokinase activity will likely result in a decrease in the levels of NADPH and GSH and an increase in NADP+ and GSSG (oxidized form of glutathione). Indeed, our results show that OGD decreases NADPH and GSH, which are significantly increased by either PARP inhibition or overexpression of HK-1 (wtHK-1 or pbmHK-1). These data demonstrate that PAR-mediated hexokinase inhibition in OGD-exposed neurons results in the depletion of an essential cellular antioxidant, GSH making the neurons more susceptible to oxidative stress. Supporting this notion, our data show that the levels of ROS are increased in OGD and significantly decreased either by PARP inhibition or HK-1 overexpression. These results indicate that PAR-mediated inhibition of HK-1 is a critical pathophysiological pathway that results in cellular bioenergetics defects and makes neurons susceptible to oxidative stress. HK (HK-1 and HK-2) is a mitochondrial-associated protein that interacts with VDAC and protects mitochondrial membranes against permeability transition and loss of Δψ_m_.^38^ Our data shows that OGD resulted in PAR-dependent inhibition of HK activity and caused the dissociation of HK from VDAC, making mitochondria susceptible to permeability transition. PARP inhibition or expressions of HK-1 (wtHK-1 or pbmHK-1) restores the Δψ_m_ in neurons.

In summary, this study shows that PAR-mediated alteration in HK-1 in neurons is a common pathological process that affects neuronal survival in brain ischemia (Figure 10). This PAR-HK-1 pathological interaction causes cellular bioenergetic defects via PAR-mediated glycolytic inhibition and subsequent impairment in mitochondrial function. This process also results in the dysregulation of oxidative homeostasis via inhibition of PPP flux, making neurons susceptible to oxidative stress while making neuronal mitochondria susceptible to permeability transition by displacing HK-1 from the mitochondrial outer membrane (Figure10) These results reveal that PARP activation and PAR accumulation is a key pathological process in brain ischemia, and pharmacological strategies targeting PAR/HK-1 interaction to restore cellular glycolysis and HK- 1 function is a potential therapeutic avenue for neuroprotection in stroke.

**FIGURE 10.**
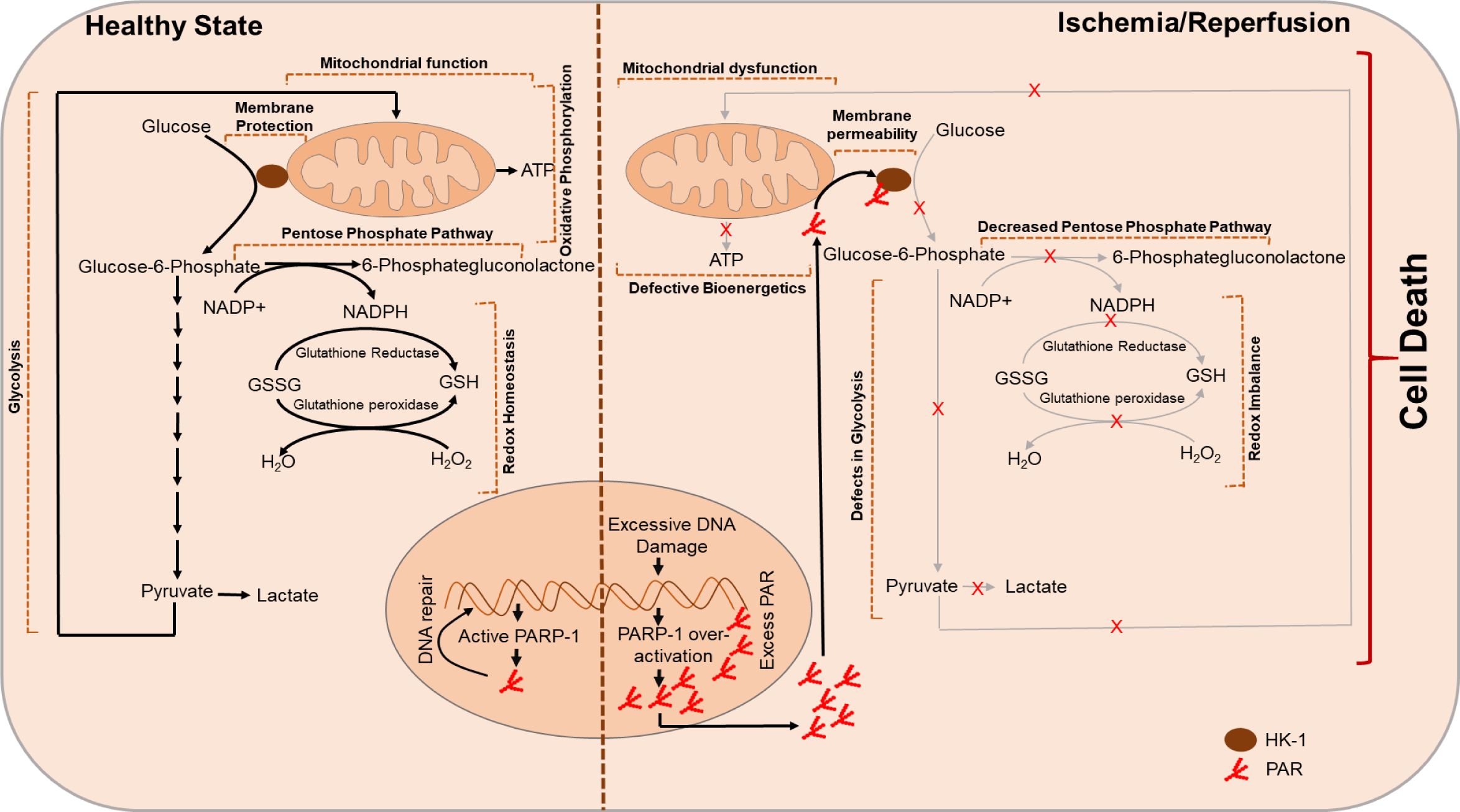
The left side of this summary figure shows that hexokinase (HK-1 in the brain) is a vital enzyme to initiate glucose metabolism by converting glucose to glucose-6-phosphate (G6P) to generate pyruvate in the glycolytic pathway. The mitochondrial Krebs cycle uses pyruvate to create substrates for the electron transport chain and oxidative phosphorylation. G6P is also utilized via the pentose phosphate pathway to maintain redox homeostasis via NADPH to convert oxidized glutathione (GSSG) back to reduced glutathione (GSH). HK-1 is localized to the outer mitochondrial membrane via interaction with VDAC. Mitochondrial localization of HK-1 is critical to protect against mitochondrial permeability transition. The right side of the figure shows that in ischemia/reperfusion, PAR binds to HK-1 and inhibits its function, resulting in bioenergetic collapse due to the loss of glycolysis and mitochondrial function. Inhibition of HK-1 by PAR also decreases the pentose phosphate pathway that causes redox imbalance and oxidative stress. PAR can also induce the dissociation of HK-1 from mitochondria rendering mitochondrial membranes susceptible to permeability transition.

## Abbreviations

PARP-1: Poly (ADP-ribose) polymerase-1;
PAR: PARP-1: Poly (ADP-ribose) polymer;
G6P: glucose-6-phosphate;
PPP: pentose phosphate pathway;
PARG: poly (ADP- ribose) glycohydrolase;
PBM: PAR-binding motif;
OGD: oxygen-glucose deprivation;
DPQ: 3,4- dihydro-5-[4-(1-piperidinyl)butoxy]-1(2H)-isoquinolinone;
MNNG: N-methyl-N’-nitro-N’- nitrosoguanidine.

## AUTHOR CONTRIBUTIONS

## ACKNOWLEDGMENTS

This work was supported by grants from the NIH R01NS08695301 (SAA), R01NS109212 (SAA), R37 NS067525 (VLD and TMD). TMD is the Leonard and Madlyn Abramson Professor in Neurodegenerative Diseases.

## DATA AVAILABILITY SATEMENT

All reported data are contained within the article. Reagents and plasmids described in this article are available upon request

